# The economic burden of protecting islands from invasive alien species

**DOI:** 10.1101/2021.12.10.471372

**Authors:** Thomas W Bodey, Elena Angulo, Alok Bang, Céline Bellard, Jean Fantle-Lepczyk, Bernd Lenzner, Anna Turbelin, Yuya Watari, Franck Courchamp

## Abstract

Biological invasions represent a key threat to island ecosystems, with pronounced impacts across environments and economies. The ecological impacts have received substantial focus, but the economic costs have lacked synthesis at spatial and temporal scales. Here we utilise the InvaCost database, the most comprehensive global assessment of published economic costs of invasive species, to assess reported spend by cost types and socioeconomic sectors, and to examine temporal trends in spending, across islands that differ in their political geography - nation states, overseas territories or offshore islands of continental countries. We based this assessment on 1473 unique cost entries comprising 2914 annual costs totalling almost US$100 million in area-corrected costs between 1965-2020. We find that offshore islands of continental countries incur the greatest total and management costs. However, nation states incurred the greatest damage costs whilst substantially financing management actions, and spent an overall greater proportion of their GDP. In contrast, spending within overseas territories was significantly lower in all respects. The most impacted sector was authorities and stakeholders, demonstrating the key role of government in addressing island invasions. Temporal trends revealed continual increases in spending across all island types. This likely reflects ongoing introduction rates globally alongside an increased recognition of the importance of islands as biodiversity hotspots, and an appetite to tackle invasive species at larger and more socially complex scales. However, the high economic costs of invasions on islands substantiates the need to prevent them in order to avoid this dire threat to biodiversity and its burden on limited conservation resources.

## Introduction

Alien species, species that have been distributed beyond their natural range through human agency, are one of the driving forces of the restructuring of global and regional species distributions and compositions (Capinha et al. 2015, Russell et al. 2017; IPBES 2019). A subset of these alien species, so termed invasive alien species (IAS), can have severe, multi-faceted impacts in their novel environments (Reaser et al. 2007, Blackburn et al. 2011). For example, they can result in reductions in ecosystem integrity e.g. loss of ecosystem services such as coastal protection or inshore productivity (Orth et al. 2006, Graham et al. 2018). IAS also create declines in local economies e.g. through crop damage or reduced availability of wild food stocks (Naylor 1996, Ballew et al. 2016). Lastly, IAS can have significant impacts on human health e.g. increased costs and hospitalisations as a result of vector borne diseases (Mwebaze et al. 2010, Mavian et al. 2019).

IAS are especially problematic in island ecosystems, which are hotspots of global biodiversity (Myers et al. 2000, Mittermeier et al. 2005) with high levels of endemism and evolutionary distinctiveness (Whittaker 2007, Kier 2009). Simultaneously, islands are among the hotspots of invasion globally, with the vast majority of the world’s islands and archipelagos supporting invasive species (Atkinson 1985, Bellard et al. 2016, 2017, Dawson et al. 2017, Turbelin et al. 2017). They have also experienced the greatest increases in alien species richness (van Kleunen et al. 2015), and IAS are one of the leading causes of native species extinctions on islands, and the most common threat after resource use i.e. harvesting or collection (Blackburn et al. 2004, Bellard et al. 2016, Doherty et al. 2016).

Increased recognition of the importance of island taxa to global biodiversity has resulted in substantial IAS management efforts on islands worldwide (Towns and Broome 2003, Veitch et al. 2011, Russell et al. 2017, Bellard et al. 2017, Veitch et al. 2019). In addition, small spatial scales make management measures feasible and useful to protect or restore biodiversity (Courchamp et al. 2003, Jones et al. 2016). Therefore, islands have proven to be the ideal testing ground for developing management strategies and deploying a range of techniques across invasion stages, from early detection (e.g. using environmental DNA [Takahara et al. 2013]), through population management of target IAS (e.g. using stable isotope techniques [Bodey et al. 2011]), to complete eradication (Towns and Broome 2003, Holmes et al. 2019). Such IAS management measures are typically costly and require significant resources, strategic planning and workforce capability. Hence, the geographic location of the islands in question, as well as their political situation, might be crucial in the success of IAS management efforts. Political and administrative differences, particularly when combined with their respective financial resources, may significantly affect potential or actual investment in IAS management. For example, the vast majority of small island states are considered to be developing nations, while in contrast, island overseas territories (OT) are exclusively administered by more developed nations (Dawson et al. 2014, Churchyard et al. 2014, Soubreyan et al. 2015, Vaas et al. 2017, Sieber et al. 2018). This discrepancy in the availability of financial resources may further enforce differences in ability to control IAS and protect native biodiversity when balanced against other urgent socio-economic and societal needs, particularly within small island developing states.

An economic rationale for addressing the impacts of IAS is often necessary for justifying action (or inaction), both for governmental and non-governmental bodies. While cost-benefit analyses have frequently been conducted prior to specific management actions or prioritisation exercises (Dawson et al. 2014, Holmes et al. 2019, Carter et al. 2021), such actions tend to be conducted on a case-by-case basis, either operationally or within a specific location. Indeed, it has been frequently argued that prevention of IAS impacts is less cost intensive than post-invasion adaptation and mitigation measures (Leung et al. 2002, Timmins and Braithwaite 2002; Russell et al. 2015). Beyond island-, or even country-specific contexts, we are currently lacking a global synthesis of economic costs of IAS on islands (Reaser et al. 2007). This is concerning as such a synthesis helps to identify knowledge and/or management gaps both locally and internationally, and highlights actions that can produce synergies across sectors. Such win-win scenarios are particularly important for nations with limited financial resources.

Using the recently developed ‘InvaCost’ database (Diagne et al. 2020), we collated reported costs of IAS on islands worldwide. Specifically, we compared reported costs of management (control, biosecurity, etc.) and damages (agricultural losses, etc.) associated with IAS, across socioeconomic sectors among islands differing in political geography (nation states, overseas territories or offshore islands of continental countries). We also compared temporal trends in spending across these different island categories. We hypothesised that i) the spending ratio between control/management and damage mitigation will be equivalent across the three island categories. That is, the economic impacts of IAS are felt, and responded to, similarly on all islands. However, as a result of their ties to larger and often financially wealthier states ii) overseas territories and offshore islands of continental countries would have proportionately greater expenditures, particularly as these locations are frequently managed for their contribution to biodiversity. Finally, in light of the ongoing increase in, and recognition of IAS impacts, we hypothesized that iii) all islands would experience increasing expenditures over time.

## Methods

### Invacost dataset

We used the InvaCost_3.0 database (Diagne et al. 2020), a publicly available living repository which compiles the reported monetary impacts of invasive species globally (https://doi.org/10.6084/m9.figshare.12668570). Diagne et al. (2020) developed InvaCost via standardized literature searches (via the Web of Science platform and the Google Scholar and Google search engines), coupled with opportunistic, targeted searches where data gaps were identified. Analogous searches were conducted in more than 10 non-English languages in e.g. French, Spanish, Chinese and Japanese (Angulo et al. 2021). Costs were extracted from the found sources and standardised to a common currency (2017 US dollars) based on annual average market exchange rate and inflation factors.

### Data Processing

#### a) Economic costs and island characteristics

Within the InvaCost database, costs were categorized by method reliability and implementation type, as well as by cost type and impacted sector, as explained below. To derive our islands database (Fig 1, Table S1), we screened the full database by the following available metadata columns:

i. First, to derive all island-associated costs, the ‘Official_country’ column was used to select all island nation states, and then column ‘Location’ was used to include all costs associated with offshore islands of continental nations, such as the Galapagos (offshore islands of Ecuador) or Reunion (OT of France).
ii. Next, to provide the most robust, conservative estimates of the costs of island invasions, we considered only highly reliable (entries assigned with *high* in the ‘Method reliability’ column) and observed (entries assigned with *observed* in the ‘Implementation’ column) costs in our estimations, thus excluding entries that were not from peer-reviewed literature or official reports (e.g., government documents), or were otherwise not reproducible (labeled as *low reliability*), as well as those that were expected but not empirically observed (labeled as *potential*, Diagne et al. 2020). However, we retained studies where costs observed at a small scale within an island were extrapolated to larger areas of the same location. We thus consider this approach, which results in 1,473 unique cost entries, to provide minimum but robust estimates of the economic impacts of IAS on islands. There is no guarantee that inclusion of low reliability data or potential costs would produce a comprehensive estimate, as many undocumented costs also undoubtedly occur (Diagne et al. 2020, 2021).

**Fig 1.**
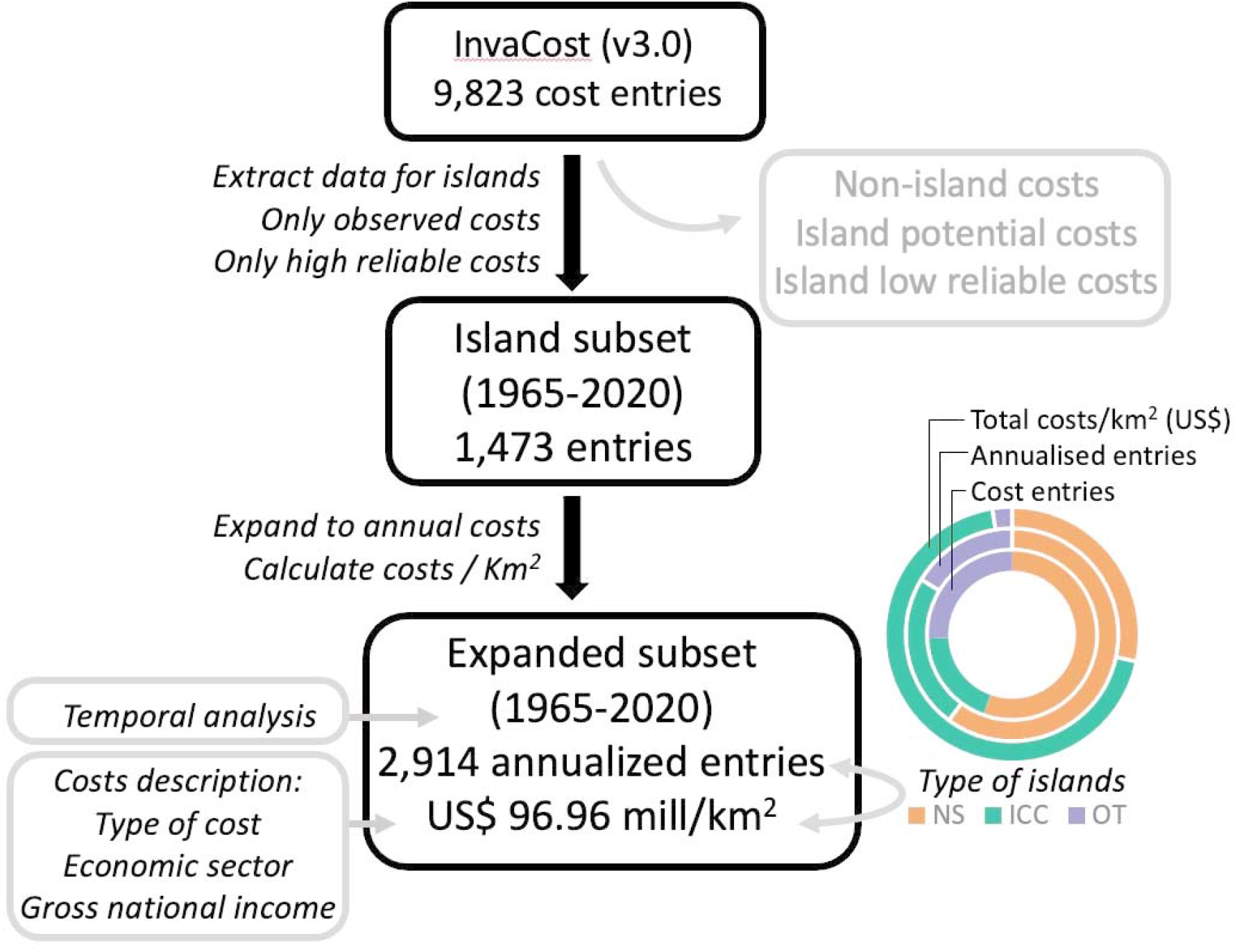
Workflow showing the extraction and filtering processes to determine the reported economic costs of invasive alien species occurring on islands. The graph depicts the proportion of entries, the proportion of annualised costs records and proportion of total costs in US$ millions km^-2^ spent by island type (ICC: Island of Continental Country, OT Overseas Territory and NS: Nation State).

Then, in order to allow for accurate comparisons, costs were standardised to costs in 2017 US$ per km^2^ using information provided by the ‘Spatial_scale’ column on the unit of measurement, when necessary returning to the primary source for clarification. Where no smaller unit of measurement was given, the cost was assumed to apply across the entire island.

Finally, we used the three categories of cost types from the ‘Type_of_cost_merged’ column: *Management* (e.g. any form of population control, biosecurity, eradication effort), *Damage* (e.g. repairs to infrastructure, human health impacts) and *Mixed* (including cost types from both previous categories and the very small quantity of unspecified costs [<0.0001%]). The ‘Impacted_sector’ column was combined into broader categories of socioeconomic impact, namely: Authorities-Stakeholders, Primary Industries (agriculture, forestry, fisheries etc), Health and Social Welfare, Environment and Mixed (including costs across multiple sectors).

#### b) Island characteristics

Costs were compared across a common metric through currency standardisation and scaling to a unit of cost km^-2^ (see above). We then added two additional classifiers: i) ‘Island Type’ to divide islands across three distinct political geographic categories: island Nation States (NSs, this includes all sovereign island nations present including small island developing states); offshore Islands of Continental Countries (ICCs); and Overseas Territories (OTs), and ii) an income classifier using the World Bank’s gross national income (GNI) assessment (four categories: lower, lower middle, upper middle, upper income country [World Bank Development Indicators]). We also added the most recent 10-year average of the relevant country’s GDP (i.e. the NS itself, or the continental country or ultimate administrative country of the island or OT in question).

#### c) Estimating Total Costs and Temporal Trends

Finally, in order to assess the full economic impact of cost entries, we annualized the cost values to consider the temporal frame in which they occurred, because the duration of reported costs could vary from a few months to years. Thus, we expanded rows within the database that corresponded to costs spanning multiple years in order that costs can be comparable and provide an accurate assessment of the economic costs of IAS through time. These multi-year costs were expanded using the ‘invacost’ package in R v4.0.3 (R Core Team 2020, Leroy et al. 2020) based on the ‘Probable_starting_year_adjusted’ and ‘Probable_ending_year_adjusted’ columns. When this information was unclear within the primary source (n = 18 entries with no clear starting year), we conservatively considered that costs occurred in only one year using the year determined from the source as the relevant year in question. This resulted in an expanded dataset of 2,914 annual costs covering the years 1965 - 2020. However, for the analysis of temporal trends we excluded costs that occurred before 1965 (n = 3) due to the small sample size, and also costs after 2017 (n = 706), due to publication lags resulting in substantially incomplete data for more recent years (median publication lag, defined as the difference between publication and impact years of the study, was 3 years). Temporal trends were assessed through implementing the summarizeCosts function of the ‘invacost’ package (Leroy et al. 2020), and were examined independently for total reported costs across all islands, across each island type separately, and for exclusive management and damage costs.

## Results

The complete islands dataset consisted of 1,473 unique cost entries, resulting in 2,914 annualised costs spanning the years 1965 - 2020 that summed to US$96.96 million in km^-2^ corrected costs. The majority of cost entries were from NSs, but the greatest economic costs were reported from ICCs (Fig 2, Table S2). Geographically, spending on islands was dominated by the Pacific region (45% of costs including both NSs and islands with political ties to Oceania, Europe, South and North America) and Europe (41% of costs, Fig 2).

**Fig 2.**
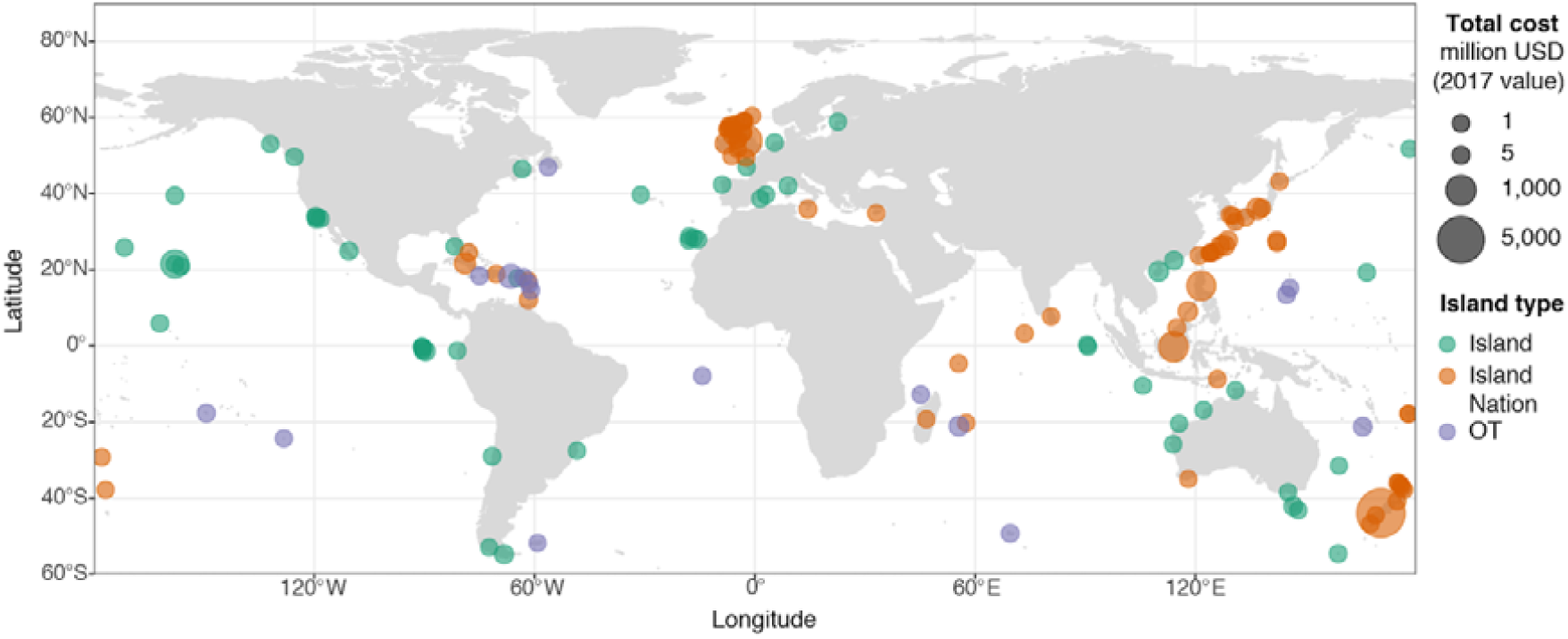
Global locations and magnitudes of reported economic costs of invasive alien species on islands. All costs are in 2017 US$ km^-2^

Across all islands, the large majority of costs (61%) were classified as exclusively management related, with exclusive damage costs comprising a much smaller proportion (3%) of the total (Fig 3a). However, the breakdown of this spending was not equivalent across island types, with ICCs spending the most on exclusive management costs and the most in total. NSs spent nearly as much on management, and incurred the majority of damage costs, although a substantial proportion of ICC costs were classified as mixed, and so contained some unknown proportion of damage within them (Fig 3a, Table S3). Total reported spending in OTs was an order of magnitude lower for management, and almost two orders of magnitude lower for damage than in the other island types. While the ratio of management to damage spending was broadly equivalent between ICCs and OTs, this decreased in NSs as a result of the higher damage spend (Table S3). However, as a proportion of GDP, there were large differences between each island type, with NSs spending the greatest fraction of their available income on IAS costs, an order of magnitude higher than the proportion of GDP spent in ICCs, and two orders of magnitude higher than the OT’s ultimate administrative country’s GDP (Fig 3b, S1, Table S3).

**Fig 3.**
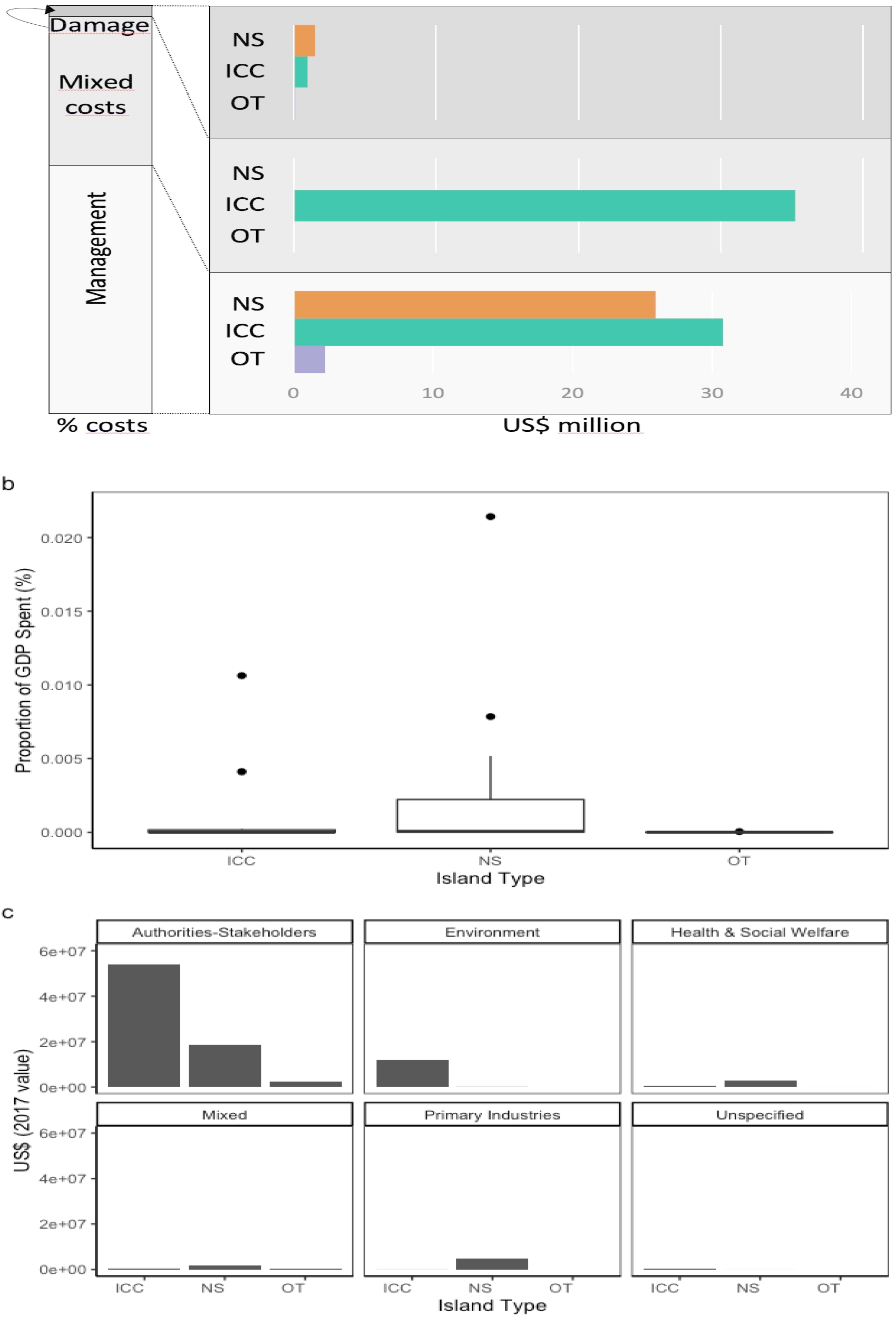
a) Distribution of costs by proportion across main cost types and by US$ millions km^-2^ across island categories b) Spend as a proportion of GDP by island type c) Distribution of costs across principal socio-economic sectors. (NS - Nation State, ICC - Island of Continental Country, OT -Overseas Territory)

Considering costs by socioeconomic sectors revealed dominant spending incurred by the Authorities-Stakeholders category (77% of annualised costs), with lesser costs attributed to the Environment (12%) and Primary Industries (5% - of which >90% comprised unspecified agricultural-forestry costs) categories (Fig 3c). Cost by sector was also not evenly spread across island types, with almost all Primary Industry costs incurred by NSs, whereas costs to Authorities-Stakeholders and the Environment dominated on ICCs. In terms of principal species incurring costs (for this we considered only costs that were clearly assigned to single species) there was almost no overlap among island types or between the principal species incurring management or damage costs (Fig 4). NS and ICC management spending was dominated by mammals and flowering plants, whereas OT management spending was dominated by insect related costs. Damage spending, although a much smaller proportion of the total, was also spread more diversely across classes in all island types (Fig 4). When compared by gross national income, the data was dominated by high and upper middle income category countries, with only a single cost estimate for a low income country, and with low and lower middle income countries representing only 0.2% of all reported annualised costs (Table S4).

**Fig 4.**
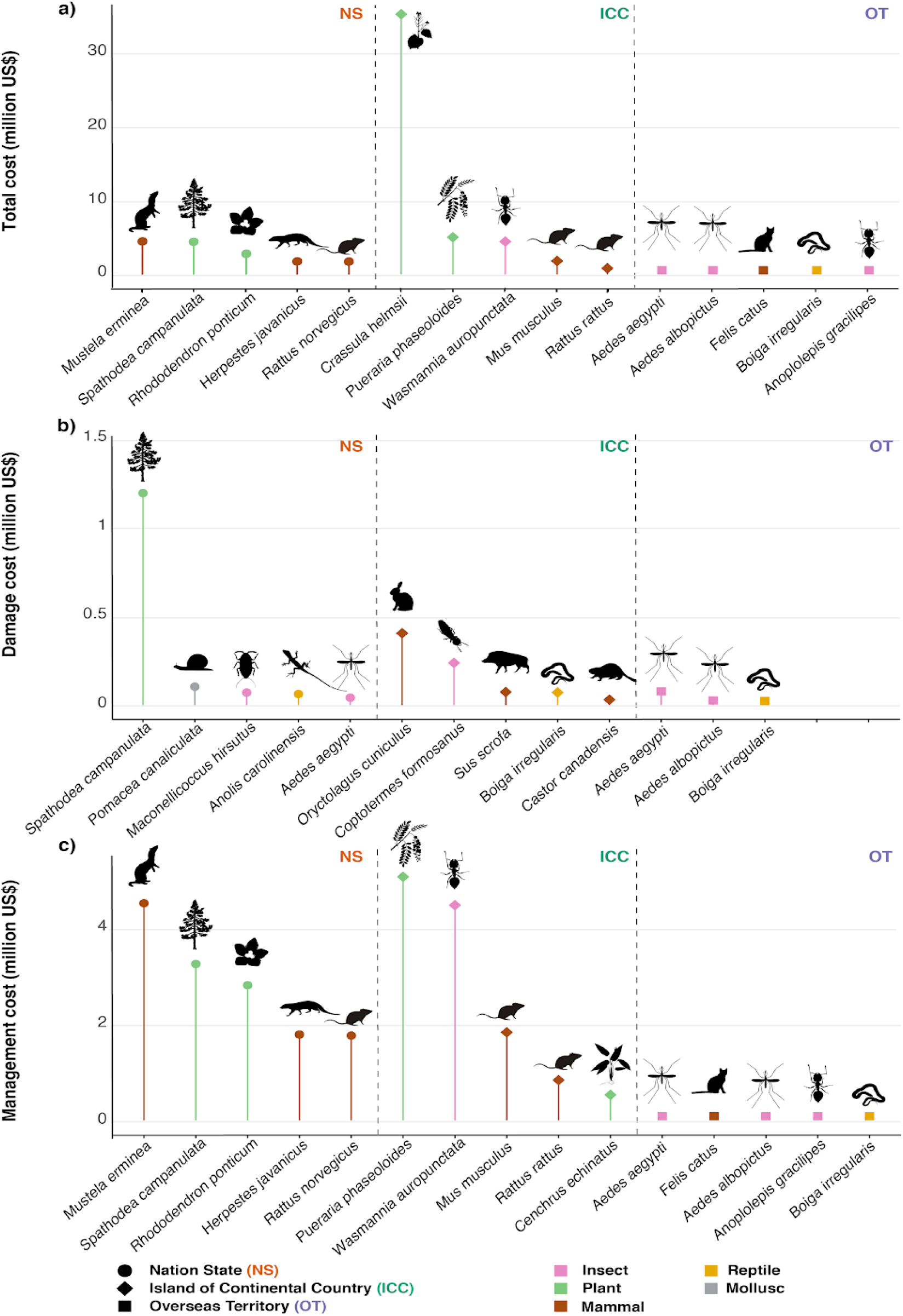
Top five most costly species by island type for (a) all costs, (b) exclusive damage costs and (c) exclusive management costs. All costs are in US$ km^-2^. There was almost no overlap between species across different island types: NS (circles), ICC (diamonds), OT (squares). Note only three species had exclusive damage costs within OTs.

When examining trends in spending through time, when all islands are considered together, there is an increasing trend in total reported costs through each decade from 1960 to 2017 apart from the 1990s, although step changes over this period are clearly evident (Fig 5). This trend is mirrored, where data is available, for all island types considered separately, with NSs exhibiting the decline in costs in the 1990s, but trends for both ICCs and OTs continually increasing. It is also true for both management and damage costs when considered separately, but with management costs consistently at least an order of magnitude greater (Fig 5).

**Fig 5.**
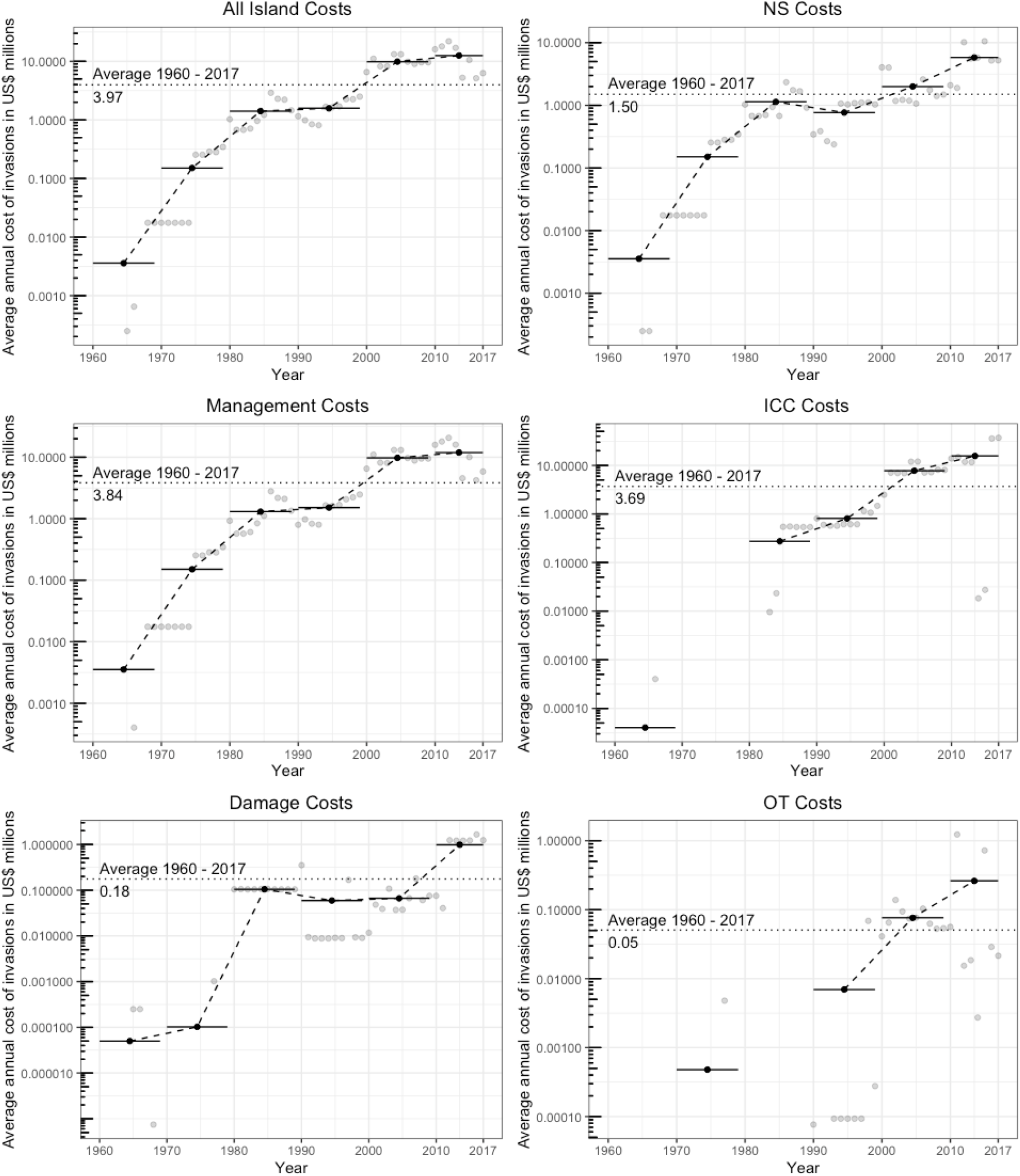
Temporal trends in reported total spending on the economic impacts of IAS across different type of costs and island types. Note the differing and logarithmic scales of the y axes when comparing expenditure.

## Discussion

We found that reported spend in insular systems to address the economic impacts of IAS comprised almost US$100 million in area-corrected costs over the past 55 years, but that there was significant variation in the types of costs across islands, particularly in respect to their political geography. Our first hypothesis - that the spending ratio between control/management and damage mitigation would be equivalent across island categories - was not supported by our results. Instead, reported ratios were lowest for Nation States (NS), with management spend far higher compared to damage in other island types. Our second hypothesis, that Overseas Territories (OT) and Islands of Continental Countries (ICC) would have greater total expenditure as a result of their connections to wealthier countries, was, surprisingly, similarly not fully supported. While ICCs did incur the greatest costs in total and over time (Figs 2, 3), this trend was not mirrored in OTs, where spending substantially lagged behind both ICCs and NSs. However, in agreement with our third hypothesis, spending across all islands combined, or across any island types considered separately, has continued to increase through time, potentially reflecting both the continuing emergence of alien species and recognition of an increasing ability to address these issues.

### Patterns of spending

Management costs dominated spending, comprising approximately two thirds of all reported expenditure on islands. However, this proportion is likely substantially higher given the large number of mixed management-damage costs reported within ICCs (Fig 2). The large proportion of spending devoted to management is in contrast to most individual countries or broader regions where costs due to IAS damage consistently outstrip management expenditure (Haubrock et al. 2021, Diagne et al. 2021 but see Bodey et al. 2021 and Watari et al. 2021). Geographically, spending was greatest in the Pacific region and western Europe, with a notable lack of reported costs from island-rich, biodiversity hotspots such as the Caribbean, SE Asia and the western Indian Ocean (Mittermeier et al. 2005). NSs incurred the greatest costs for damages, with lower to negligible figures reported by ICCs and OTs respectively. It is notable that NSs spent a far greater (although still extremely small) proportion of their GDP on economic costs from IAS. This may reflect the greater impact of IAS on NS economies where, for example, damage to crops or other primary industries directly impacts the country’s capacity to feed itself or increase its export capacity (Reaser et al. 2007, Mwebaze et al. 2010). Indeed, reported costs to primary industries occur almost exclusively within NSs, where there may be a greater recognition of the need to protect finite and limited resources. Furthermore, NSs may also have an increased recognition of the value of their unique biodiversity for generating income from natural capital or ecotourism (Fotiou et al. 2002, Dolins et al. 2010, Ballesteros-Meija et al. 2021). Similarly, the limited damage costs reported in ICCs and OTs likely reflect their relatively small contribution to the governing country’s overall economy (industry or infrastructure). This, coupled with small proportions of national populations resident in ICCs and OTs and their spatial isolation, and thus limited electoral influence, may influence the extent of spending reported, particularly for damage costs, for these island types.

In contrast, costs on ICCs were substantially devoted to management, with more than a 30-fold difference compared to their damage costs. ICCs are much more likely to comprise uninhabited locations where significant recordable economic damages are far less likely, particularly to socioeconomic sectors such as primary industries or human health. Rather, damage costs in these locations may reflect losses to biodiversity and ecosystem function, but placing monetary values on such losses is complex, particularly where human livelihoods are negligibly affected by impacts to ecosystem services (Nunes et al. 2001, Kallis et al. 2013, Roberts et al. 2018). On the other hand, ICCs may receive significant management spending on IAS control and eradication as they are likely targets for ecological restoration, particularly through the removal of invasive terrestrial species, following early conservation models established in New Zealand (Towns and Broome 2003, Jones et al. 2016, Veitch et al. 2011, 2019). Under this model, these islands can be used as ‘arks’ and either sustain relict and subsequently recovering populations, or receive translocations, of highly endangered naïve island endemics that may receive exceptional financial support due to their greater extinction risk (Jones et al. 1995, Carthey & Banks 2014, Russell et al. 2015). This is demonstrated by the dominance of reported costs from ICCs with highly successful IAS eradication campaigns including: Macquarie Island, Australia (Helmstedt et al. 2016), the California Channel Islands, USA (Parkes et al. 2010), the Galapagos, Ecuador (Carrion et al. 2011) and a large number of islets and islands around Mexico (Samaniego-Herrera et al. 2018). In addition, many uninhabited offshore islands are also protected areas of varying legislative status, which may increase the likelihood of IAS management in these locales. Notably, the socioeconomic sector incurring the greatest proportion of reported costs across all islands, but particularly incurred by ICCs, is that of authorities and other stakeholders, evidence that these management costs tend to fall to governmental organizations and conservation NGOs. However, it was also apparent that there was almost no consistency in the species generating the majority of either management or damage costs across island types. While there is the potential for this to reflect a lack of uniformity in the reporting of costs incurred by specific species, and no doubt also reflects some regional, latitudinal or societal differences, it also strongly suggests there is a need for greater information sharing to synergise experience across locales considering the fact that many islands support similar IAS (McKinney and Lockwood 1999, Simberloff et al. 2013).

### Under-reported or underfunded?

Regardless of the type of cost, it is clear that spending in OTs is substantially lower than on other islands, and this is seen not just in absolute cash terms, but exceedingly so in the proportion of the ultimate administrative countries’ GDP (Table 2). As these costs are corrected for area, this is not simply a reflection of the smaller landmass OTs occupy globally. Instead, it appears that countries maintaining OTs either do not record, or do not invest in, IAS management or damage mitigation at any significant scale, as there is no evidence to suggest that IAS presence is decreasing in these locations contrary to the global trend (Dawson et al. 2017, Seebens et al. 2017). Indeed, the effort expended on recording IAS costs in OTs appears to vary substantially by location. For example, New Caledonia represents the highest number of IAS cost entries, though not the highest spending, within the French *départements*, with Réunion also recording substantial numbers of IAS and associated costs (Renault et al. 2021). However, simultaneously, there is little data on IAS presence or costs for other French OTs including Mayotte, Martinique and French Guiana (Turbellin et al. 2017, Renault et al. 2021). In contrast, Cuthbert et al. (2021) do not include IAS costs from OTs in an examination of the impacts of IAS to the UK economy, potentially as a result of limited reporting reflecting a general pattern of oversight and undervaluing of these locations (Churchyard et al. 2014). Most OTs represent small islands with limited internal resources comparable to many small island nations amongst the NSs, and without substantial support from the ultimate governing country, they are likely to struggle to conserve the biodiversity they support (Churchyard et al. 2014, Key 2017, Vaas et al. 2017). While the InvaCost database collected data on monetary values, and so may fail to capture some costs of biodiversity or cultural losses, for example the ecosystem functions of extinct island endemics (Zavaleta et al. 2001, Wood et al. 2017), it is unlikely that the data collection process missed so much spending in OTs that it would then be equivalent to other island types, especially when such values may also go unrecorded elsewhere.

Our work also highlights the almost complete lack of cost records from developing nations, with >99% of all reported island costs occurring in countries classified by the World Bank as higher or upper middle income countries. Given that several of the largely or completely absent countries represent mega-diverse biological hotspots (Mittermeier et al. 2005), this highlights a pressing research or assessment need or, minimally, an urgent effort to increase the ability to produce and disseminate publications (Wallace et al. 2020). The absence of reported costs may reflect a true knowledge gap that, in turn, may reflect a difference in perceived priorities between countries that have pressing needs around sectors such as human health and education. It may also reflect a limited number of studies due to poorer funding opportunities as compared to their richer counterparts. In either event, there is an urgent need to improve our understanding of the costs of IAS in these locations due to the likely severe negative impacts invasive species have on a range of indicators, including health, productivity, happiness, child poverty levels, and declines in ecosystem services (United Nations 2017) as well as biodiversity (Simberloff et al. 2013).

### Temporal trends

Given the importance of islands to global diversity, temporal trends in spending on management exhibit an encouraging trend across all islands where, regardless of political geography, average decadal spending has increased almost uniformly between 1960 and 2017. In addition, this expenditure on management has consistently been at least an order of magnitude greater than that spent on damage (Fig 3). While publication delays mean that estimates for the most recent years are necessarily an underestimate, this trend is certain to continue for at least the 2010s, and is in stark contrast to the repartition of reported spending on IAS globally (Diagne et al. 2020). While the specific reasons for continued spending may differ across islands of differing political geography, this potentially reflects ongoing increases in rates of introductions (Seebens et al. 2017, Lenzner et al. 2020), allied to increased recognition of the importance of islands as biodiversity hotspots (Mittermeier et al. 2005) and an increasing appetite to tackle IAS at larger and more socially complex scales (Oppel et al. 2011, Russell et al. 2015, Veitch et al. 2019, Carter et al. 2021). Such efforts have the potential to lead to substantial conservation gains (Jones et al. 2016, Holmes et al. 2019). However, translating approaches from uninhabited to inhabited islands is challenging, and the logistical difficulties, social challenges and necessarily increased costs associated with such endeavours may constrain the capacity or willingness of governments to commit to such endeavours given the current higher risk of failure (Oppel et al. 2011, Russell et al. 2015, Harper et al. 2020).

### Conclusion

Using the most comprehensive available data on the economic impacts of IAS on islands, we demonstrated that the political geography of an island is central to the type and quantity of expenditure and/or the likelihood of reporting costs. Nation States were more likely to report losses due to damage caused by IAS, likely reflecting the more significant impact such losses make on their overall economies, but also revealing probable knowledge gaps as to the extent of many socio-economic impacts of IAS across all locations (*cf* Crystal-Ornelas and Lockwood 2020). There is also a clear disparity in reporting of costs from developing nations and, despite islands from all categories supporting unique biodiversity, spending was especially low in Overseas Territories. Given the importance of Overseas Territories to total country biodiversity, a greater focus of attention on the impacts of IAS in these locations would ensure European countries, in particular, made substantial progress towards the achievement of Aichi Biodiversity targets under the Convention on Biological Diversity (CBD 2011). Such an approach would benefit all islands given the strong overlap between the Aichi biodiversity targets and sustainable development goals (Schultz et al. 2016, UN 2017). Nevertheless, the predominance of spending on management approaches (incorporating all stages from biosecurity through to long-term control/eradication), and the continuing increase in this spend over time across all islands, suggests that there continues to be the potential to make substantial conservation and development gains in insular ecosystems. In particular, the high economic burden imposed by biological invasions on islands adds to the strong evidence that prevention rather than management will better protect insular biodiversity.

## Declarations

### Availability of Data and Material

The analyses are based on a subset of the InvaCost Database which is available at Figshare https://doi.org/10.6084/m9.figshare.12668570 In addition, the subset used is available as Supplementary Table 1.

### Code Availability

N/A

### Authors Contributions

TWB, CB, BL & FC conceived the study; TWB, EA, CB, JFL, BL, AT & YW collected additional data, with all analyses performed by TWB, EA and AT. TWB led the drafting of the manuscript, with all authors contributing critically to drafts and giving final approval of the submission for publication.

### Ethics Approval

N/A

### Consent to Participate

N/A

### Consent for Publication

All authors have read the final draft and approve its submission for publication

## Acknowledgements

This work was conducted following a workshop funded by the AXA Research Fund Chair of Invasion Biology and is part of the AlienScenario project funded by BiodivERsA and Belmont-Forum call 2018 on biodiversity scenarios. The authors also acknowledge the French National Research Agency (ANR-14-CE02-0021) and the BNP-Paribas Foundation Climate Initiative for funding the InvaCost project and enabled the construction of the database. TWB acknowledges funding from the European Union’s Horizon 2020 research and innovation programme Marie Skłodowska-Curie fellowship (Grant No. 747120). JFL would like to thank the Auburn University School of Forestry and Wildlife Sciences for travel support to attend the InvaCost workshop. Funding for EA comes from the AXA Research Fund Chair of Invasion Biology of the University of Paris Saclay. BL acknowledges funding by the BiodivERsA-Belmont Forum Project “Alien Scenarios” (FWF project no: I 4011-B32).

