## Supplemental Tables 2-4 for "The economic burden of protecting islands from invasive alien species"

| **Island Type** | **Cost Entries** (%) | **Annualised Cost Records** (%) | **Number of Countries** | **Total Costs km^-2^**  (US$ millions) |
| --- | --- | --- | --- | --- |
| Nation State (NS) | 817  (55) | 1753    (60) | 14 | 27.5 |
| Island of Continental Country (ICC) | 277  (19) | 686  (24) | 22 | 67.0 |
| Overseas Territory (OT) | 379  (26) | 475    (16) | 4 | 2.4 |

Table S2. Breakdown of database records from InvaCost 3.0 by island type. Note that for OTs the Number of Countries refers to the ultimate administrative country rather than the location.

| **Island Type** | **Management Costs** | **Damage Costs** | | | | **Mixed Costs** | | **Total**  **Costs** | | **Mean Spend as Proportion of GDP** | | **Management to Damage Ratio** |
| --- | --- | --- | --- | --- | --- | --- | --- | --- | --- | --- | --- | --- |
| Nation State (NS) | 25.96 | | | 1.55 | | 0.02 | | 27.53 | | 0.01 | | 16.88 |
| Island of Continental Country (ICC) | 30.78 | | | 0.99 | | 35.26 | | 67.03 | | 0.001 | | 31.59 |
| Overseas Territory (OT) | 2.30 | | | 0.11 | | 0.01 | | 2.42 | | 0.00002 | | 27.35 |

Table S3. Economic costs of IAS across categories of expenditure, mean total spending as a proportion of GDP of reporting countries, and the ratio of management:damage spending within the three island types. All cost values are in 2017 US$ millions km^-2^. Mixed costs refer to those categorized as mixed or unspecified (the latter totalling <US$10000) in the data.

| **World Bank Classifier** | **Total Costs (US$ millions km^-2^)** | **Number of Cost Entries** | **Countries Represented** |
| --- | --- | --- | --- |
| Higher income country | 77.8 | 2290 | Antigua, Australia, Bahamas, Brunei, Canada, Chile, Cyprus, Estonia, France, Ireland, Japan, Malta, Netherlands, New Zealand, Portugal, Seychelles, Spain, UK, USA |
| Upper middle income country | 19.0 | 544 | Argentina, Brazil, China, Cuba, Dominican Republic, Ecuador, Fiji, Grenada, Maldives, Mauritius, Mexico, Sri Lanka |
| Lower middle income country | 0.2 | 79 | Indonesia, Philippines, Timor-l’Este |
| Lower income country | 0.005 | 1 | Madagascar |

Table S4. Economic costs of IAS in 2017 US$ millions corrected for area across World Bank classifiers of countries’ Gross National Income.
